# High Content Screening, a reliable system for *Coxiella burnetii* isolation from clinical samples

**DOI:** 10.1101/2019.12.17.880484

**Authors:** Rania Francis, Maxime Mioulane, Marion Le Bideau, Marie-Charlotte Mati, Pierre-Edouard Fournier, Didier Raoult, Jacques Yaacoub Bou Khalil, Bernard La Scola

## Abstract

Q fever, caused by *Coxiella burnetii*, is a worldwide zoonotic disease that may cause severe forms in humans and requires a specific and prolonged antibiotic treatment. Although the current serological and molecular detection tools enable a reliable diagnosis of the disease, culture of *C. burnetii* strains is mandatory to evaluate their antibiotic susceptibility and sequence their genome in order to optimize patient management and epidemiological studies. However, cultivating this fastidious microorganism is difficult and restricted to reference centers as it requires biosafety-level 3 laboratories and relies on cell culture performed by experienced technicians. In addition, the culture yield is low, which results in a small number of isolates being available. In this work, we developed a novel high content screening (HCS) isolation strategy based on optimized high-throughput cell culture and automated microscopic detection of infected cells with specifically-designed algorithms targeting cytopathic effects. This method was more efficient than the shell-vial assay when applied to both frozen specimens (7 isolates recovered by HCS only, sensitivity 91% *vs* 78% for shell-vial) and fresh samples (1 additional isolate using HCS, sensitivity 7% *vs* 5% for shell-vial). In addition, detecting positive cultures by an automated microscope reduced the need for expertise and saved 24% of technician working time. Application of HCS to antibiotic susceptibility testing of 12 strains demonstrated that it was as efficient as the standard procedure that combines shell-vial culture and quantitative PCR. Overall, this high-throughput HCS system paves the way to the development of improved cell culture isolation of human viruses.

## Introduction

*Coxiella burnetii* is the causal agent of Q fever, a polymorphic disease that may occur as acute, mostly mild and self-limiting forms, or potentially severe persistent focalized infections, the main presentations being endocarditis and vascular infections (1),(2),(3),(4),(5). Although a large number of animals are able to carry *C. burnetii*, the main reservoirs of the bacterium include sheep, goats and cattle (6). Q fever may cause outbreaks(7),(8), the largest to date having been registered in the Netherlands (9). The laboratory diagnosis of *C. burnetii* infections relies mainly on serology and molecular biology (10). Such tools have greatly improved the diagnosis and management of patients, especially those developing a persistent focalized infection such as blood culture-negative endocarditis (1). Molecular detection assays can notably detect *C. burnetii* in clinical samples before seroconversion occurs. However, these methods cannot overcome the need for culture. Culturing *C. burnetii* is restricted to reference laboratories, as this bacterium requires cell culture, is classified as a risk group 3 microorganism and thus must be manipulated in biosafety level 3 laboratories. Moreover, it is highly contagious and can be infectious at the unit level. For these reasons, cultured strains remain scarce, limiting the access to antibiotic susceptibility tests, modern whole genome sequencing for epidemiological studies and research on virulence (11). However, the genome availability of isolated strains allowed the development of more efficient molecular detection tools (12). Therefore, isolating more strains remains crucial and many strategies were developed over time to cultivate *C. burnetii* (13). Nowadays, co-culture remains the key tool for isolation. Reference centers and diagnostic laboratories have adopted the shell-vial assay for the co-culture of intracellular bacteria (14). *C. burnetii* was typically cultured in shell vials on HEL cells and detection was monitored every 10 days by immunofluorescence, Gimenez staining and specific PCR (15),(14). However, this strategy remains subjective, tedious, time consuming, operator dependent and has poor yield (R. Francis *et al.*, submitted for publication).

In this work, we revisited the isolation strategy of *C. burnetii* and brought improvements at two main axis: co-culture and detection. We started by standardizing the co-culture process of susceptible cell lines at many levels such as culture medium, temperature monitoring, contamination control, cell viability and proliferation monitoring. The detection process was then optimized using a fully automated system for high content screening that was used in a previous study for the detection of giant viruses in protozoa (16). This new generation microscope allowed the live monitoring of co-cultures and large scale image analysis, where specific algorithms were applied to detect any potential signs of infection including cytopathic effects, morphological modifications and vacuoles induced by *C. burnetii* and predict cell phenotypes.

After validating the proof of concept, a large scale comparative screening of clinical samples from patients with acute or chronic Q fever was then performed with both conventional shell-vial and high content screening strategies for *C. burnetii* isolation. Finally, this strategy was adapted for antibiotic susceptibility testing. This new strategy showed higher efficiency and sensitivity than the shell-vial assay for the isolation of *C. burnetii* from clinical samples, with easier and quick manipulations associated with reduced subjectivity and thus need for highly experienced technicians.

## Materials and Methods

In the developmental stage, we targeted two main axis: the co-culture process and the detection process. At the axis of co-culture, modifications took place at many levels: cell lines culture, microplates, cell concentrations, proliferation monitoring and the co-culture process. At the axis of detection, improvements were made at the levels of cell staining, screening protocol and data analysis to detect cytopathic effects induced by *C. burnetii* (vacuoles or cell burst), and finally, automation was introduced.

### 1. Co-culture standardization

#### a. Cell lines selection

Two cell lines were used as cellular supports for co-culture: the human embryonic lung fibroblast MRC5 cells (RD-Biotech, Besançon, France) and the mouse fibroblast L929 cells (ATCC® CCL-1). The cell lines were cultured to confluence at 37°C under 5 % CO2 in MEM (Minimal Essential Medium) supplemented with 2 mM L-glutamine per liter and 10 % or 4 % heat inactivated FBS (Fetal Bovine Serum) for MRC5 and L929 cells, respectively (15). Cells were then harvested using a phenol red free MEM culture medium, and transferred into 96 well microplates at a volume of 200 µl per well and incubated for 24 hours (h) to allow cell adhesion.

#### b. Choice of microplates

Different 96 well microplates were compared using the two cell lines cited above. The objective was to choose appropriate plates with the best compromise between image resolution, cell adhesion and confluence, as well as cell viability. We tested clear transparent plates with thick polymer bottom (Ref. 167008, Thermo Scientific), black plates with optical-bottom and coverglass base (Ref. 164588, Thermo Scientific), and black plates with optical-bottom and polymer base (Ref. 165305, Thermo Scientific).

#### c. Optimal cell concentrations

Different cell concentrations (10^5^, 2×10^5^, 4×10^5^ and 10^6^ cells/ml) were tested to determine the optimal concentration granting a confluent monolayer suitable for inoculation and monitoring. Cell viability and growth were monitored for 30 days, with the culture medium being changed every 10 days for MRC5 cells and every 7 days for L929 cells.

#### d. Cell proliferation monitoring

L929 cells have an uncontrolled growth, which could interfere with the visualization of cytopathic effects. Therefore, different attempts were made to control their proliferation and maintain a single monolayer for 30 days. However, MRC5 cells have a contact inhibition of proliferation, and therefore, no further controls were required.

##### d.1. Culture medium composition

The first attempt to control L929 cell growth was by reducing the percentage of FBS added to the culture medium. Cell proliferation was monitored using a culture medium supplemented with 2 % and 4 % FBS, respectively.

##### d.2. Cycloheximide addition

Another attempt was the addition of cycloheximide to the L929 cell monolayer, which inhibits the synthesis of nucleic acids and proteins of eukaryotic cells (17). Different concentrations, ranging from 0.05 to 1 µg/ml, were tested. Cell viability and proliferation rate were monitored for 30 days, and cycloheximide was added after every culture medium renewal. The effect of cycloheximide on *C. burnetii* infectivity and cell susceptibility to infection was assessed by quantitative PCR, Gimenez staining and immunofluorescence, to search for any inhibition or improved infection related to cycloheximide. We compared results from cells infected with *C. burnetii* with and without cycloheximide addition to the co-culture.

#### e. Coxiella burnetii strains

Three strains of *C. burnetii* were used for infection: *C. burnetii* strain Nine Mile phase II (CB NMII), *C. burnetii* strain 223 (CB 223) and *C. burnetii* strain 227 (CB 227). All strains used in this study were obtained from CSUR-IHU (Méditerranée Infection, Marseille, France). Bacterial cells were produced on L929 cells, and quantification was performed by endpoint titration (TCID50) on L929 and MRC5 cells, 15 days post infection.

#### f. Co-culture process

We kept the same co-culture strategy used in the shell-vial assay for the isolation of *C. burnetii* (14), and we optimized it at many levels. Figure 1 summarizes the steps of the isolation strategy in the conventional shell-vial assay and the new developed High Content Screening (HCS) assay. Regarding the co-culture process, cells were transferred into 96 well microplates and incubated for 24 h prior to infection. Supernatant was then removed and infection was carried out with 50 µl of CB NMII, CB 223 and CB 227 diluted up 10^-10^. Similarly to the shell vial assay, low-speed centrifugation of plates (700 x g for 1 h at 22°C) was performed to enhance the attachment and the penetration of the bacteria inside the cells. The final volume was adjusted to 250 µl with culture medium. Uninfected cells were kept as a negative control.

**Figure 1:**
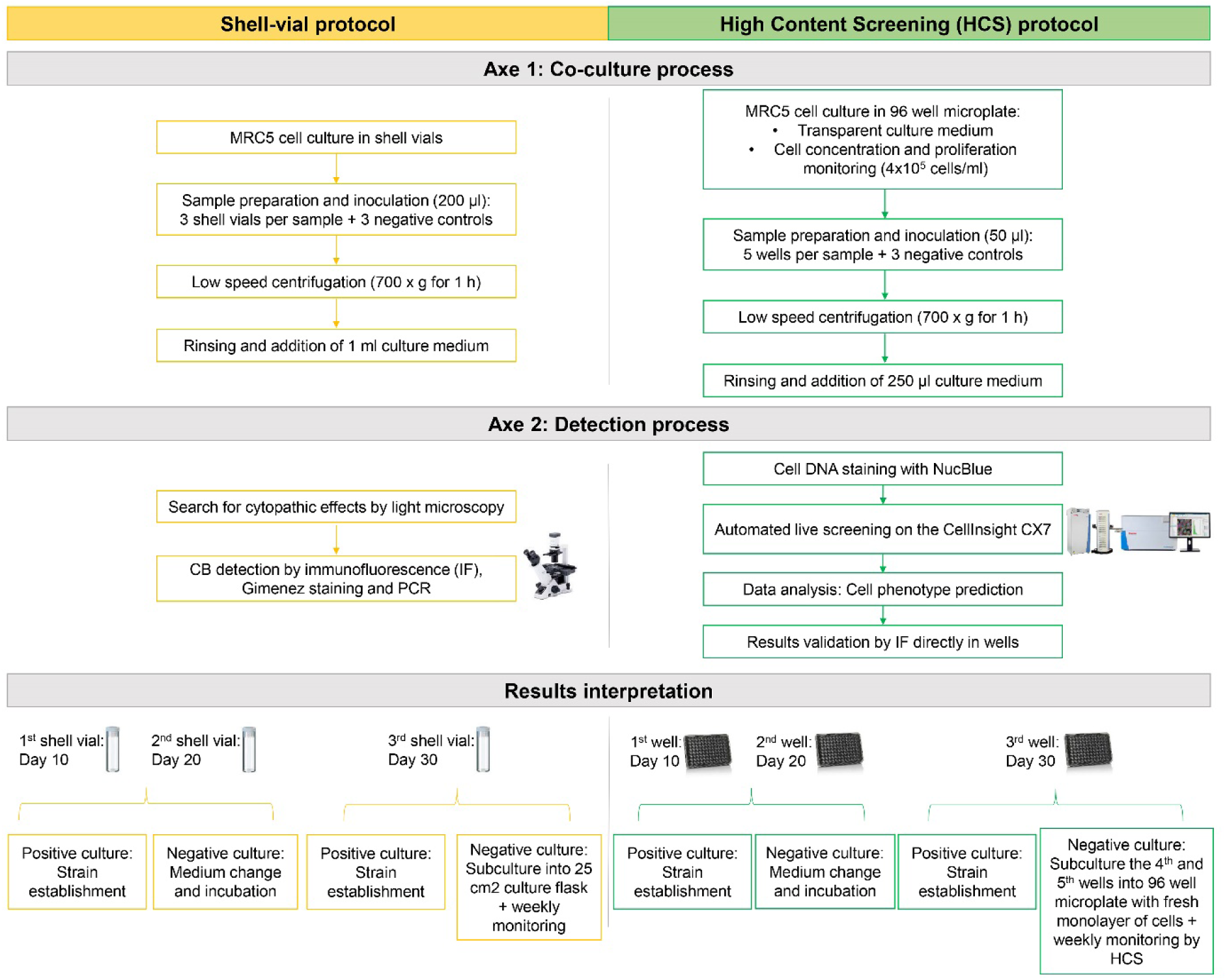
Traditional and improved co-culture and detection processes adopted for the isolation of *C. burnetii*. This figure details all the steps followed in the processing of a clinical sample using the conventional shell-vial assay and the new optimized High Content Screening technique.

### 2. Detection process optimization

The workflow for the detection process development is summarized in figure 2.

**Figure 2:**
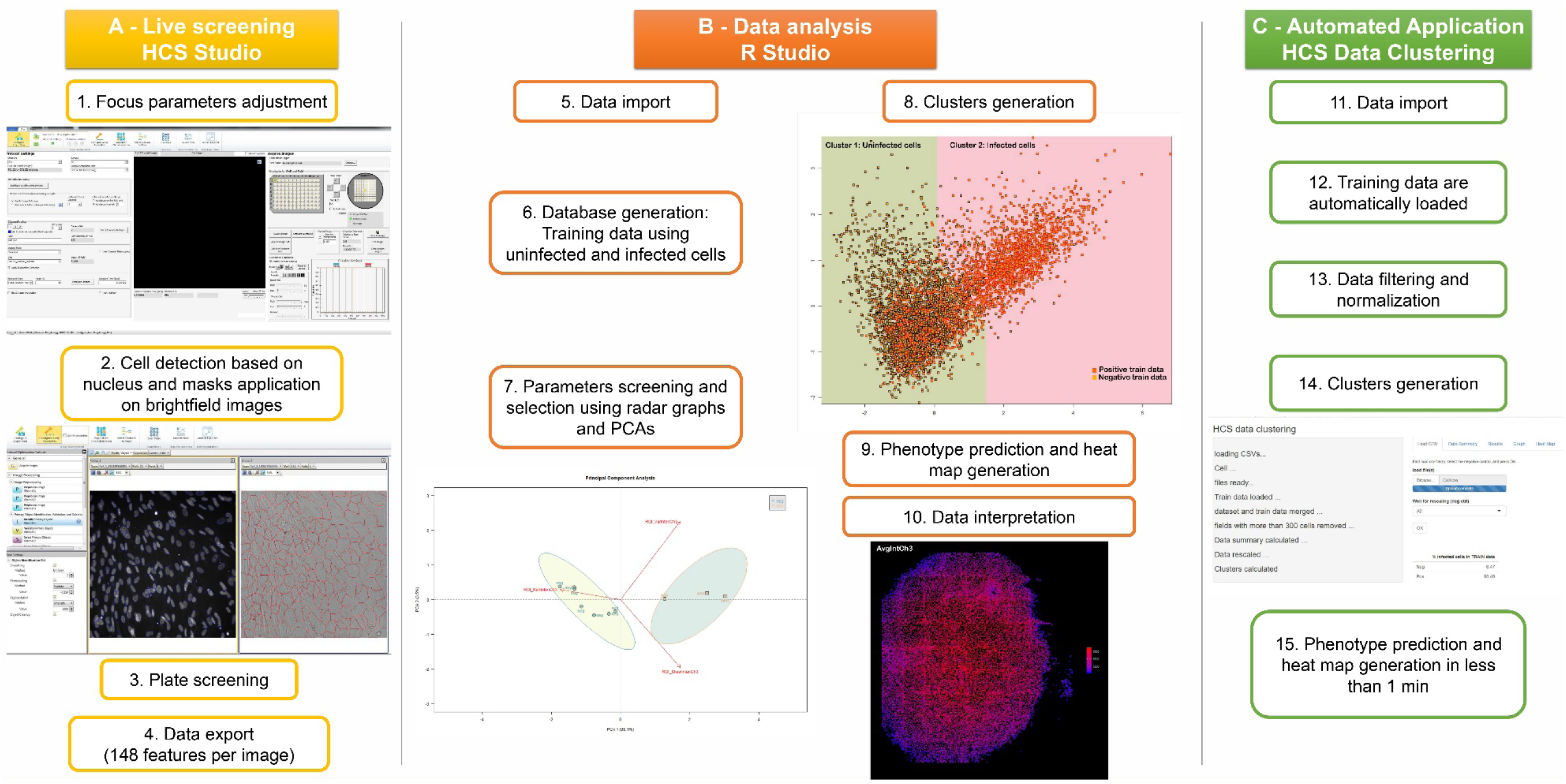
Workflow of the detection process development and optimization at the software level. (A) describes the steps performed in the HCS Studio software for image and data generation. (B) summarizes the main steps in the data analysis script development in R Studio. (C) describes the minimized application for automated analysis.

#### a. DNA staining of cells

NucBlue™ Live ReadyProbes™ reagent (Molecular Probes, Life Technologies, USA) was used as a live cell DNA stain. Staining was performed by direct addition of NucBlue to the cells without washing. Different concentrations were tested and the minimal concentration granting sufficient staining and lowest cell toxicity was adopted for each cell line. Cell viability and aspects were monitored for any cytotoxic effects related to staining.

#### b. Screening protocol

Image acquisition and analysis were performed at 10, 20 and 30 days post infection using the automated CellInsight™ CX7 High Content Analysis Platform (Thermo Scientific) allowing real time acquisition and on-the-fly multiparametric analysis. Acquisition parameters were defined in the HCS Studio 3.1 software (Thermo Scientific) using the Morphology Explorer Bio-Application. This later provides quantitative measurement of morphological and texture related features at the single cell level, intracellular level and multi-cellular level. Autofocus parameters and exposure times were adjusted so that the fluorescent or optical density signal reached 50 % the dynamic range of the 14-bit camera. The nuclear fluorescent probe NucBlue (386 nm) was used to perform software-based autofocus and served as a primary mask for single cell detection and quantification. The secondary channels consisted of brightfield images with adapted z offset to collect morphological information. The region of interest (ROI) consisted of an enlarged Voronoi diagram derived from the nuclear mask to include the cytoplasm, thus limiting the possible overlapping with neighboring cells. Cell aggregates and debris were excluded from analysis using area cutoffs. Image acquisition was performed using a 20x objective (0.45 NA) and 80 images or fields were generated per well in a way to cover 90% of the well surface. We extracted intensity, texture and morphology related information from the region of interest and exported a dataset of 148 features for every cell.

#### c. Data analysis

Developed algorithms described in this paper were performed on MRC5 cells infected with 3 strains of *C. burnetii*: CB NMII, CB 223 and CB 227. Cells infected with *C. burnetii* exhibit a particular phenotype due to vacuoles formation. Therefore, we used the extracted dataset to detect cytopathic effects and differentiate between infected and uninfected cells. The exported file was uploaded in a dedicated application developed in R Studio® using the user interface ShinyR. We generated a database of labeled cell data coming from uninfected and infected cells. Outliers were removed from this dataset and 6000 cells were kept as training data. Using radar graphs and Principal Component Analysis (PCA), we screened all 148 features and identified key features that distinguish infected cells (positive control) and uninfected cells (negative control). Two K-Means clusters, representing infected and uninfected cells respectively, were calculated using the generated training data and key features. These clusters were then used to predict the phenotype of the experimental dataset using a semi-supervised K-Mean clustering algorithm (form flexclust R package, kccaFamily “k-median”). A preliminary data sorting was performed based on the total cell count per well in order to detect wells showing cell burst and prevent false negative results. All wells with less than 5000 cells per well were excluded from analysis and marked as “not applicable” (NA) in the final result. The prediction algorithm was then applied to the remaining dataset. This algorithm associates a phenotype for each cell depending on the cluster it falls in, and therefore, the percentages of infected and uninfected cells per well can be calculated. Finally, we defined a threshold for positivity based on the percentage of infected cells per well.

#### d. System automation

After validation, we coupled the CellInsight™ CX7 microscope with an automation system consisting of a robotized incubator Cytomat™ 2C-LIN (ThermoScientific) and a plate handler Orbitor™ RS Microplate mover (Thermo Scientific). Incubation times, plate handling and acquisition protocols were monitored through the Thermo Scientific™ Momentum 5.0.5 software. The script for data analysis was also automated using the user interface ShinyR in order to make the application user friendly and interactive. An automated application, HCS Data Clustering, was created where all the steps (cell data import, data filtering and normalization, clustering of training data, prediction of experimental data and results generation) were automatically performed.

### 3. Comparison of the detection process to the Gold standard methods

The immunofluorescence assay and the manual quantification of infected cells were adopted as reference methods in order to validate the results.

#### a. Immunofluorescence assay

We kept the immunofluorescence assay as a gold standard method for results validation. The same protocol previously described for the detection of *C. burnetii* (14) was optimized in microplates. Imaging was performed on the CellInsight™ CX7 microscope, and the entire well was screened at 20x magnification.

#### b. Manual quantification of infected cells

We then performed a manual quantification of infected cells using brightfield images. We quantified the number of vacuoles visible to the naked eye and then calculated the percentage of infected cells per well for each strain as follows: Percentage of infected cells = (Number of vacuoles / Total cell count) x 100. Results were then compared to the HCS results.

### 4. Proof of concept validation: Artificial samples

In order to test the system’s efficacy towards clinical samples, we artificially contaminated one blood sample and one serum sample with CB NMII at different concentrations (Pure, 10^-3^ and 10^-6^ dilutions). The selected samples had negative PCR results for *C. burnetii*. Co-culture was then performed on MRC5 cell line as described above. Fifty µl were used for inoculation, and cells were rinsed twice with culture medium after the centrifugation step. Two negative controls were considered: uninfected cells and cells co-cultured with the non-contaminated samples. Co-cultures were then monitored every 10 days on the automated CellInsight™ CX7 microscope and results were validated by immunofluorescence.

### 5. Applicative stage: High Content Screening assay versus shell-vial assay

#### a. Screening of clinical samples

For the applicative stage, we performed a comparative study on 90 clinical samples from our samples collection using the traditional shell-vial assay technique (14) and the new optimized HCS technique. A large variety of samples was tested including blood, valves, biopsies, thrombi, aneurysms, abscesses, sera, articular fluids and tick samples. The first group consisted of 47 frozen samples from which different strains of *C. burnetii* were previously isolated after primary inoculation upon reception at the laboratory. The second group consisted of 43 fresh samples tested positive by PCR as previously described (12) and inoculated prospectively. Co-cultures were performed on MRC5 cell line simultaneously in shell vials as previously described (14) and in microplates for HCS. Briefly, the shell-vial assay consists of inoculating 200 µl of the sample onto a monolayer of cells. 3 shell vials were inoculated for each sample and then centrifuged at 700 x g for 1 h at 22°C. Cells were then rinsed twice with PBS (phosphate-buffered saline) and incubated at 37°C with 1 ml of culture medium. 3 shell vials containing uninfected cells were kept as negative control. Co-cultures were monitored under a light microscope for cytopathic effects detection after 10, 20 and 30 days post infection. The results were validated by immunofluorescence, Gimenez staining and specific PCR (15),(14). Subcultures were performed as previously described by Raoult *et al.* (14). As for the HCS strategy, 5 wells were inoculated for each sample at a volume of 50 µl per well. The plates were then centrifuged and the cells were rinsed. The final volume was then adjusted to 250 µl with culture medium. 3 wells containing uninfected cells were kept as negative control. Monitoring was performed at the same time points on the CellInsight™ CX7 microscope, and the results were validated by immunofluorescence. After 30 days, negative co-cultures were sub-cultured into 96 well microplates containing a fresh monolayer of cells and then monitored weekly using the same strategy. We then compared the results from both strategies regarding isolation rate and culture delay.

#### b. Application for Minimal Inhibitory Concentration (MIC) testing

We adopted the same principle developed by Angelakis *et al.* for antimicrobial susceptibility testing. We tested the MIC of two antibiotics used in the treatment of *C. burnetii*: doxycycline and hydroxychloroquine (18), using both the conventional shell-vial strategy and new HCS technique. 12 strains of *C. burnetii* were tested: CB 109S, CB 196, CB 226, CB 228, CB 242, CB 244A, CB 248, CB 249A, CB 250, CB 252, CB 260 and CB Henzerling. Regarding the HCS strategy, strains were cultured in 96 well microplates containing a monolayer of MRC5 cells with serial two-fold dilutions of doxycycline (0.25-8 µg/ml) and hydroxychloroquine (0.25-4 µg/ml). Uninfected cells treated and not with the highest antibiotic concentrations were used as negative controls and the positive control consisted of infected cells without any antibiotic treatment. Each test was performed in quadruplicate and results were assessed 15 days post infection by HCS for cytopathic effects detection. In parallel, the standard MIC testing was performed in shell vials and results were assessed by quantitative PCR as previously described (18).

### 6. Statistical analysis

The R Studio® software was used to perform all statistical tests included in strategy development and data analysis.

### 7. Ethical statement

According to the procedures of the French Commission for Data Protection (Commission Nationale de l’Informatique et des Libertés), collected data were anonymized. The study was approved by the local ethics committee of IHU (Institut Hospitalo-Universitaire) - Méditerranée Infection.

## Results

### 1. Co-culture standardization

#### a. Culture medium and microplates selection

The use of a transparent culture medium without phenol red indicator minimized the auto-fluorescence that could interfere with the imaging process. All 96 well microplates tested showed adequate cell adhesion, however, plates with coverglass base were less suitable for prolonged culture durations (Fig. S1). On the other hand, black plates with optical-bottom were better for imaging than clear plates with thick polymer bottom, as photo-bleaching was minimal and a better resolution was obtained, especially on brightfield images. Therefore, we adopted the black plates with optical-bottom and polymer base to be used for co-culture.

#### b. Optimal cell concentrations

Different cell concentrations were tested for each cell line to determine an optimal concentration granting a confluent monolayer for 30 days without cell overgrowth. The optimal concentrations were at 4×10^5^ cells/ml and 2×10^5^ cells/ml for MRC5 and L929 cells respectively (Figure 3).

**Figure 3:**
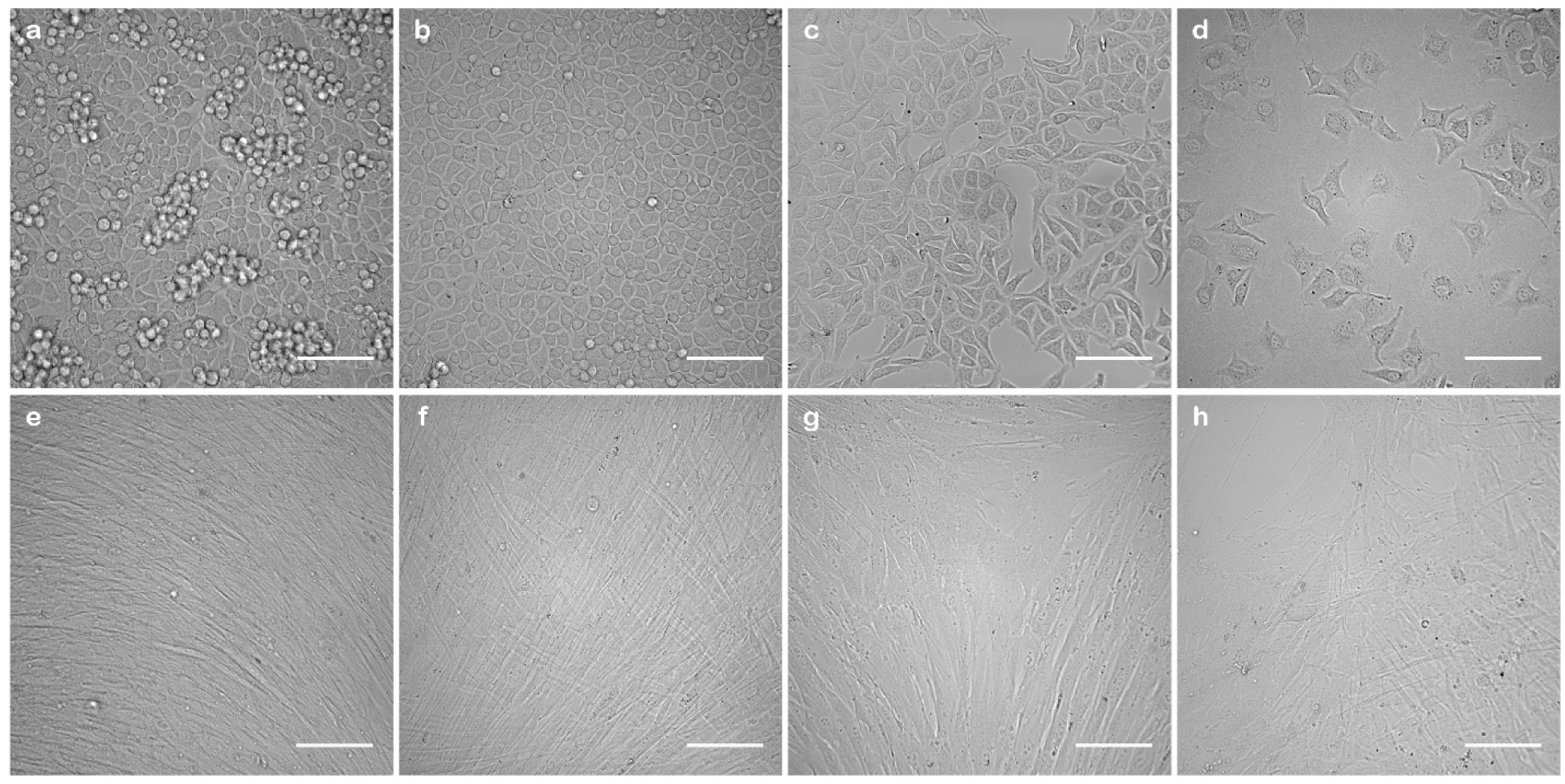
Cell density of L929 and MRC5 cell lines at different initial concentrations, 24 h into culture. (a), (b), (c) and (d) represent respective brightfield images of L929 cells at 10^6^, 4×10^5^, 2×10^5^ and 10^5^ cells/ml. (e), (f), (g) and (h) represent respective concentrations for MRC5 cells. Scale bars indicate 100 µm.

#### c. Cell proliferation monitoring

Contrary to MRC5 cells that have contact inhibition of proliferation, L929 cells showed uncontrolled cell proliferation and aggregates started forming 3 days into culture (Fig. S2 – d, e, f). Moreover, when infected with *C. burnetii*, cytopathic effects or vacuoles were difficult to visualize due to high cell density (Fig. S4 - b). Therefore, controlling cell overgrowth was a must to maintain a single monolayer of cells for the longest period.

##### c.1. Culture medium composition

We started by reducing the percentage of FBS added to the culture medium from 4 % to 2% to check if cell proliferation would be slower. However, no significant change in the proliferation rate was observed and cells became very dense starting 3 days into culture (Fig. S3).

##### c.2. Cycloheximide addition

We tested a wide range of cycloheximide concentrations and searched for cytotoxicity or cell mortality, while monitoring the proliferation rate. High concentrations showed extensive cytotoxic effects on cells and induced rapid cell mortality. The optimal concentration was 0.25 µg/ml for an initial cell concentration of 2×10^5^ cells/ml. It is important to note that cycloheximide was only added to cells after the 24 h period allowing cell adhesion. However, although proliferation rate was lower, it was not completely inhibited, and therefore, we found it necessary to increase the cycloheximide concentration up to 0.5 µg/ml after culture medium renewal at days 7, 14 and 21. This strategy allowed us to maintain a single monolayer of L929 cells for 30 days with no significant toxicity or mortality (Fig. S4).

Moreover, no significant difference in infectivity was observed between cells treated or not with cycloheximide in terms of *C. burnetii* infectivity and/or L929 cell susceptibility to infection (Fig S4 and Fig. S5). Immunofluorescence and Gimenez images showed similar infection states in cells treated or not with cycloheximide. The same was observed by PCR quantification, where the bacterial multiplication rate was the same. However, cytopathic effects visualization was not possible in the absence of cycloheximide, where high cell density masked the vacuoles formed by *C. burnetii* (Fig. S4 – b, c).

### 2. Optimized detection process

#### a. Cell DNA staining

Regarding DNA staining, several concentrations of NucBlue were tested and optimal concentrations granting sufficient staining were 4 ng/ml and 2 ng/ml respectively for MRC5 and L929 cells for the pre-determined cell concentrations. This corresponds respectively to 10 and 5 µl per well added directly from the stock solution. Note that NucBlue is a live cell stain and was directly added to culture without cell wash. Cell aspects and viability were monitored by microscopy to search for any cytotoxicity related to staining. We noticed that prolonged contact with cells induced nuclear fragmentation, morphological modifications, and eventually, cell mortality (Fig. S6). To overcome this problem, staining was performed a few hours before screening and stained wells were only considered exploitable during the following 24 h.

#### b. Screening protocol

Image acquisition and analysis protocol was developed in the HCS Studio software to extract the maximum data from the region of interest in brightfield images. MRC5 cells infected with serial dilutions of CB NMII, CB 223 and CB 227 were then screened at 10, 20 and 30 days post infection using the CellInsight™ CX7 microscope. Screening time was found to be around 3 minutes per well, where autofocus, image acquisition and algorithms application on generated images were simultaneously performed. 80 fields were screened per well and 4 images were generated in each field, where the first consisted of the nucleus fluorescence image, followed by 2 brightfield images and the overlay image. Cell data containing intensity, texture and morphology information were extracted from generated images as a .csv file and used in the following step of the analysis.

#### c. Data analysis

Cell data extracted from MRC5 cells infected with CB NMII, CB 223 and CB 227 were used for data analysis and database generation. A database of negative and positive controls was generated to be used as training data. We selected data from different time points of infection. Positive controls were selected from wells where ∼50 % of cells were infected. Highly infected cultures are often in advanced states of cell death and do not resemble early stages of infection, thus, it was better to use data from images with moderate infection rate (∼50 %) and to find a clustering that meets this value while leaving the negative train data as close to zero as possible. We then identified 4 key features that distinguish well between the negative and the positive controls using radar graphs and principal component analysis: nuclear average fluorescence intensity per cell, skewness or the levels of asymmetry of the brightfield intensity distribution around the mean within the region of interest, kurtosis or the levels of peakedness or flatness of the brightfield intensity distribution within the region of interest and finally the ratio of the variation intensity over the average intensity of the brightfield within the region of interest (ObjectAvgIntenCh1, ROI_SkewIntenCh3, ROI_KurtIntenCh3 and Var_Avg.IntensityRatio respectively). The later feature was calculated to compensate for the loss of illumination at the well edges, a phenomenon known as the vignette effect, observed in ROI_VarIntensityCh3 (the standard deviation of intensities in the region of interest). Training data normality was assessed in a QQ plot. Data were then filtered accordingly and outliers were removed to ensure a normal distribution. Using the 4 key features and the training data, 2 clusters were generated using the K-Means clustering algorithm. These clusters represent uninfected and infected cells, respectively. Due to experimental variability, training data were rescaled so the mean values of negative training data equal the mean values of untreated cells for each feature (untreated cells being the negative control of the experimental dataset to be predicted). The clustering algorithm predicted a baseline of 6 to 7 % infected cells in the negative training data and a value of 50 to 60 % infected cells in the positive training data. These values are the agnostic result generated by the clustering algorithm. The predicted baseline of infected cells in the negative control is due to the presence of artifacts, such as debris or dead cells, which could interfere with the results.

Therefore, any prediction below this baseline was considered as a negative result. We empirically determined a threshold of 10 % for positivity. However, any data between 7 and 10 % were systematically checked for possible false negativity. The percentage of infected and uninfected cells per well of the experimental dataset was then predicted using the generated clusters. Results were then represented as a color-coded heat map showing the percentage of infected cells (Figure 4-A). All values below the baseline of 7 % are represented in white (negative result), values between 7 and 10 % are represented in green (suspected), and values above 10 % are represented in red (positive result). However, wells showing cell burst are prone to false negative results, therefore, wells with less than 5000 cells were excluded from analysis and marked as NA in the heat map (gray color).

**Figure 4:**
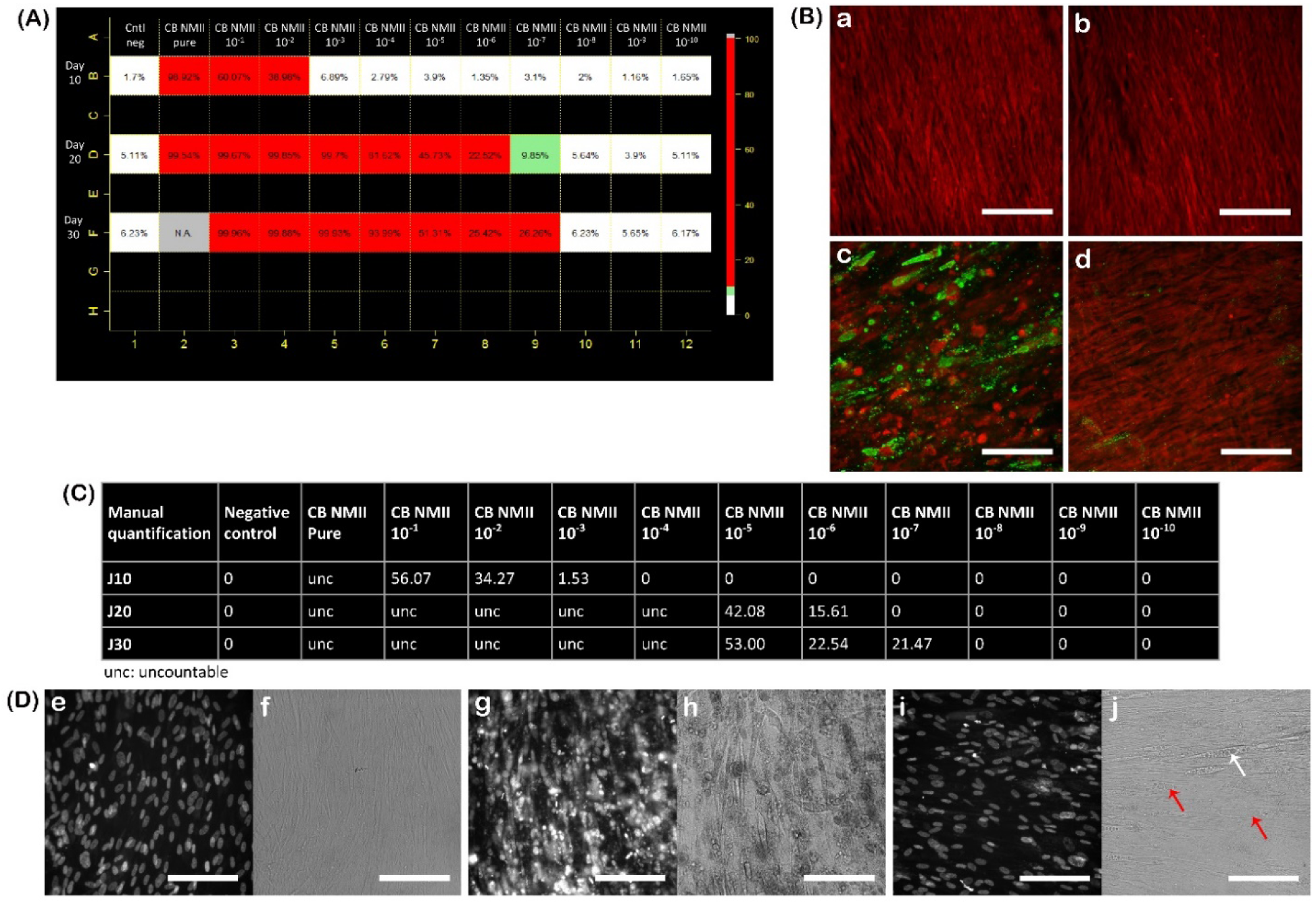
Results from the prediction algorithm and validation references of MRC5 cells infected with CB NMII at 10, 20 and 30 days post infection. (A): The heat map represents the percentages of infected cells obtained with the prediction algorithm of MRC5 cells infected with serial dilutions of CB NMII. (C): The table represents the manual quantification results from the same experiment. (B): Respective immunofluorescence images of (a) the negative control (well B1 in the heat map), (b) the false positive result (well D9), (c) a positive result (well B3) and (d) a false negative result (well B5). (D): Respective fluorescence and brightfield images of (e, f) the negative control, (g, h) cells at an advanced stage of infection and (i, j) slightly infected cells (vacuoles are indicated with red arrows). Scale bars represent 100 µm.

#### d. Automated system

The automation system allowed the systematic handling of several plates, where plates are transported by the robotic arm from the incubator to the microscope and vice versa (https://www.mediterranee-infection.com/acces-ressources/donnees-pour-articles/plate-handler-orbitor-rs-microplate-mover). Momentum software supervised the incubation time of each plate and synchronized the appropriate acquisition protocols developed in HCS Studio software. This fully automated system allows minimal handling of plates by the operator and thus reduces the risk of cross contamination. Regarding data analysis, all steps were automatically performed in the automated application HCS Data Clustering. The time required for analysis was less than 1 min from data import to results generation.

### 3. Detection process validation

Plates infected with 3 strains of *C. burnetii* were used in the developmental stage for algorithm optimization. Prediction results were generated as color-coded heat maps and were validated by immunofluorescence as well as by a manual quantification of infected cells (Figure 4 and Table S1). Figure 4-A shows an example of the generated heat map of MRC5 cells infected with CB NMII at 10, 20 and 30 days post infection. Regarding manual quantification, the number of vacuoles visible to the naked eye was quantified on brightfield images and the percentage of infected cells was then calculated. Wells showing advanced stages of infection were difficult to quantify and were noted as uncountable (unc). Note that manual quantification was very difficult to perform and was highly time consuming. We obtained values close to the predicted percentages, however, not as accurate due to the difficulties in quantification. However, manual quantification helped detecting false positive and false negative results. We detected 0.68 % false negative results and 7.08 % false positive results. False negative results were predicted below the 7 % baseline, and further investigation showed a very low infection rate, where only a few vacuoles were detected on the generated images and by immunofluorescence. Furthermore, false positive results were predicted as suspected or positive (> 7%) when no vacuoles were detectable. False positivity can be due to several reasons like high cell density, dead cells or debris… Table S1 summarizes the results of the prediction algorithm by HCS versus the manual quantification for each strain. Different infection profiles were observed for each strain, where the infection was positive up to 10^-7^ dilution for CB NMII, 10^-5^ dilution for CB 223 and 10^-6^ dilution for CB 227, 30 days post infection.

**Table 1:** Minimal Inhibitory Concentrations of doxycycline and hydroxychloroquine for different *C. burnetii* strains tested by the conventional qPCR technique and the new HCS strategy.

| *Coxiella burnetii* strains | MIC doxycycline qPCR | MIC doxycycline HCS | MIC hydroxychloroquine qPCR | MIC hydroxychloroquine HCS |
| --- | --- | --- | --- | --- |
| **CB 109S** | 0.25 µg/ml | 0.25 µg/ml | >4 µg/ml | >4 µg/ml |
| **CB 196** | 0.25 µg/ml | 0.25 µg/ml | >4 µg/ml | >4 µg/ml |
| **CB 226** | 0.25 µg/ml | 0.25 µg/ml | 4 µg/ml | 4 µg/ml |
| **CB 228** | 0.25 µg/ml | 0.25 µg/ml | >4 µg/ml | >4 µg/ml |
| **CB 242** | 0.25 µg/ml | 0.25 µg/ml | 4 µg/ml | 4 µg/ml |
| **CB 244A** | 0.25 µg/ml | 0.25 µg/ml | >4 µg/ml | >4 µg/ml |
| **CB 248** | 0.25 µg/ml | 0.25 µg/ml | >4 µg/ml | >4 µg/ml |
| **CB 249A** | 0.25 µg/ml | 0.25 µg/ml | >4 µg/ml | >4 µg/ml |
| **CB 250** | 0.25 µg/ml | 0.25 µg/ml | >4 µg/ml | >4 µg/ml |
| **CB 252** | 0.25 µg/ml | 0.25 µg/ml | >4 µg/ml | >4 µg/ml |
| **CB 260** | 0.25 µg/ml | 0.25 µg/ml | >4 µg/ml | >4 µg/ml |
| **CB Henzerling** | 4 µg/ml | 4 µg/ml | >4 µg/ml | >4 µg/ml |

### 4. Proof of concept validation: artificial samples

The prediction algorithm was as efficient with clinical samples as with the pure bacterial culture in the detection of cytopathic effects. However, more false positive results were observed due to debris coming from samples (Fig. S7).

### 5. Applicative stage: High Content Screening assay versus shell-vial assay

#### a. Screening of clinical samples

Among the first group of 47 frozen samples from which *C. burnetii* was previously isolated, isolation ratios were of 37/47 (78.7%) for the conventional shell-vial assay and 43/47 (91.5 %) for the HCS assay. Regarding the second group of 43 prospectively inoculated specimens, isolation ratios were of 2/43 (4.7 %) and 3/43 (7 %) respectively. Results are shown in figure 5 and Table S2. Overall results for the conventional shell-vial and assay and the HCS assay were of 39/90 (43 %) and 46/90 (51 %). The majority of the isolated strains originated from valves samples. Moreover, 32 % of the strains were isolated faster with the HCS than the shell-vial technique, 28 % were isolated at the same time points with both techniques and only 23 % were isolated faster with the shell-vial technique. Note that 95 % of strains isolated with the HCS assay were recovered before 30 days post infection, compared to 65 % with the shell-vial assay. These results show significantly higher efficiency and isolation rate of the new HCS strategy compared to conventional methods, where 7 strains were recovered from different clinical samples solely using the HCS assay. Furthermore, we compared the operating time required for each step of the process with both strategies on 20 clinical samples with 5 % positivity rate; We observed more time consumption (24%) during manipulations with the shell-vial assay (25 h) than the HCS assay (19 h) (Table S3)

**Figure 5:**
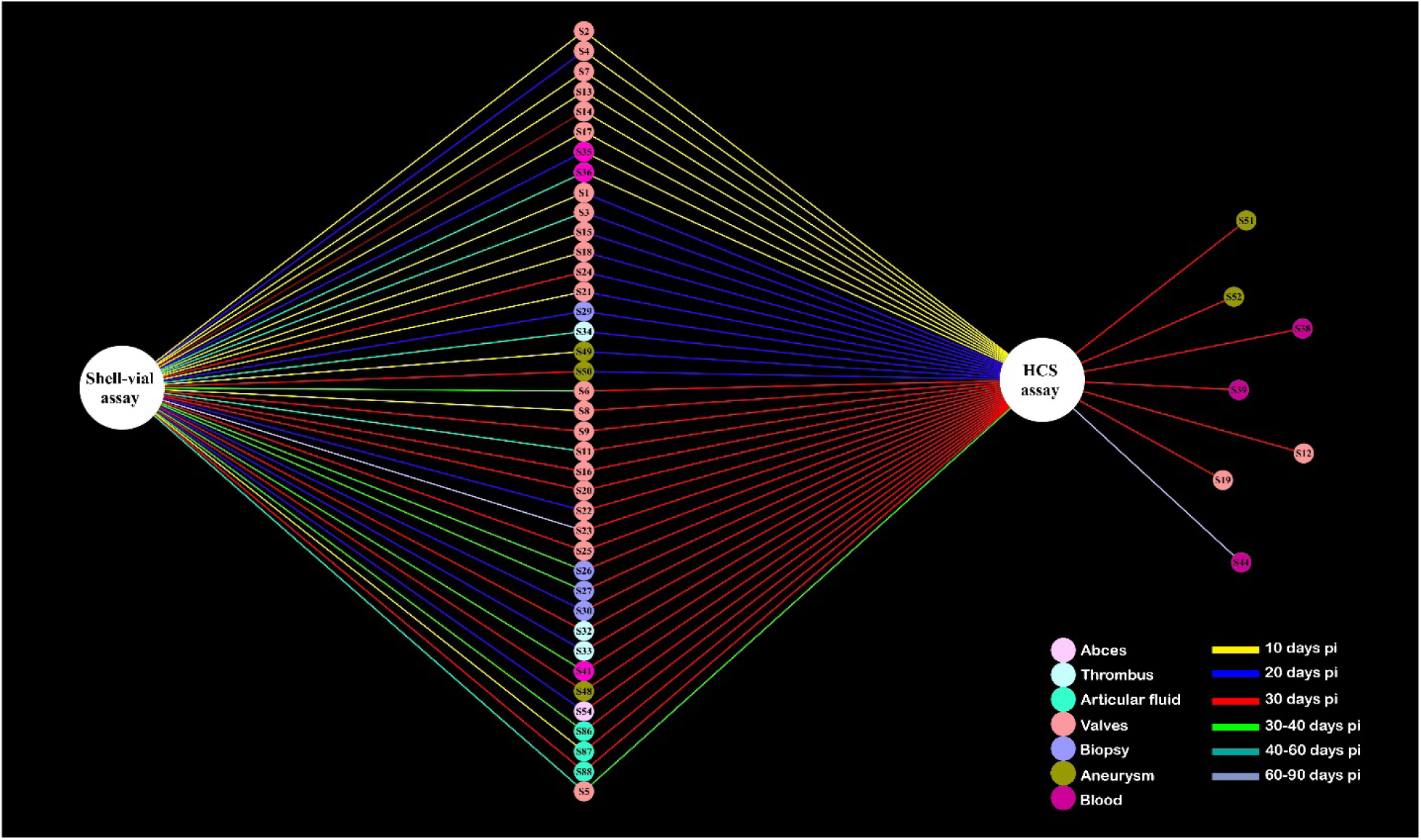
Comparative results between the shell-vial assay and the HCS assay for the isolation of *C. burnetii* from clinical samples. This interaction scheme shows all positive samples isolated using the two strategies, the sample’s origin (nodes) and their co-culture delay (lines).

#### b. Application for Minimal Inhibitory Concentration (MIC) testing

The results of MIC testing of 12 *C. burnetii* strains are summarized in table 1. Similar results were obtained in both cases, where HCS strategy detected cytopathic effects due to *C. burnetii* multiplication and the conventional technique quantified bacterial multiplication by quantitative PCR.

## Discussion

Over the past decades, major questions regarding intracellular bacteria like *C. burnetii* started to be resolved after the isolation and the proper identification of strains (1),(3),(19),(20). Currently, the rapid diagnosis of Q fever is possible with various culture-independent tools such as serology, molecular biology and histology (10),(21),(22). However, culturing the bacterium remains crucial as assessing its infectivity, tropism and virulence may only be obtained from isolates (23). Recently, many attempts were made for the axenic culture of *C. burnetii* on agar plates or in cell-free liquid medium (24),(25). Although this approach was successfully used for primo-isolation and to propagate established strains, cell culture currently remains the reference method for isolation (14),(15). Therefore, updating and improving cell culture by introducing novel technologies is mandatory. For this, we developed a new isolation strategy starting from culture standardization to the detection process optimization through an automated imaging platform and data analysis for cell phenotype prediction.

We observed that uncontrolled cells can affect susceptibility to infection and complicate the detection of the pathogen. The fact that cells are not controlled, the multilayers will mask the detection of cytopathic effects or vacuoles. Many samples were found to be false negative, where PCR results were positive but no signs of infection were detectable by microscopy. This is common and usually associated with poor sample transport and conservation, susceptibility of the bacterium and previous antibiotic therapy. Therefore, monitoring cell concentrations and proliferation, as well as choosing an adequate culture medium are critical factors for a more efficient co-culture. It is also important to avoid temperature fluctuations which can stress the cells and cause derivation, i.e. cancer cell lines are not the best choice for optimal culture, and thus primary cell lines should be used in the future. In addition, the use of microplates instead of shell vials for co-culture had many advantages, where co-culture and immunofluorescence are performed in sealed microplates which protects the culture as well as the manipulator from contamination. Another advantage was realizing the immunofluorescence in wells as we can overlay results with brightfield images which is more quantitative and less risky than shell vials. Moreover, managing samples in microplates is better where we can culture up to 18 samples in a single plate, which is equivalent to 57 shell vials, and thus manipulations are easier and incubators are less crowded.

Regarding the axis of detection, scanning in microplates didn’t change the area of screening corresponding to the same as in shell vials. In addition, we noticed that screening the whole well by the robotic microscope is always sure and explores the totality of the surface, whereas observation of the shell vial under an inverted microscope is more difficult, requires expertise and covers only a small part of the surface with less resolution. Therefore, our new HCS system showed a higher isolation rate in reduced time points compared to the conventional shell-vial assay, in addition to higher sensitivity and specificity, as well as reduced subjectivity. It is important to note that the choice of samples was dependent on their availability and we managed to isolate *C. burnetii* even though samples were frozen. A small rate of false negative results (0.68 %) was observed in cases where infection was very low, and false positive results (7.08 %) depended on the cell status and the amount of debris present in the well. Nevertheless, this risk is easily corrected by immunofluorescence and specific PCR in suspected samples.

The introduction of a panel of cell lines for the isolation of *C. burnetii* may increase the system’s efficiency to isolate more strains having different susceptibilities. Previous studies have already described the variation of the susceptibility of *C. burnetii* to different cell lines, as well as strain dependent susceptibility (26),(27). This system would also be applicable for tropism and virulence assessment.

Using a panel of cell lines for isolation would easy with this automated system that allows the processing of several plates at the same time under incubation with reduced cross contamination risks. In addition, this method does not require any expertise besides performing co-culture, since the screening process and results extraction are automated, and any biologist, student or technician can manage to recover the data.

We successfully applied this new strategy to study the antimicrobial susceptibility of *C. burnetii* which can replace the conventional PCR technique since this method is more feasible, economic, and faster than PCR. In addition, PCR results do not always reflect the number of infectious particles.

Finally, this high content screening method was based on semi-supervised deep learning and algorithms are subject to being updated and optimized for different applications regarding other intracellular bacteria as well as viruses.

## Acknowledgments

This work was supported by a grant from the French State managed by the National Research Agency under the “Investissements d’avenir (Investments for the Future)” program with the reference ANR-10-IAHU-03 (Méditerranée Infection) and by the Région Provence-Alpes-Côte-d’Azur and the European funding FEDER PRIMI.

We sincerely thank all Thermo Fischer team for their technical support on the High Content Analysis platform and the automation system.

The authors declare that Maxime Mioulane is an employee at Thermo Fisher Scientific.

